# Bio4j: a high-performance cloud-enabled graph-based data platform

**DOI:** 10.1101/016758

**Authors:** Pablo Pareja-Tobes, Raquel Tobes, Marina Manrique, Eduardo Pareja, Eduardo Pareja-Tobes

**Affiliations:** oh no sequences! research group, Era7 bioinformatics

## Abstract

**Background:** Next Generation Sequencing and other high-throughput technologies have brought a revolution to the bioinformatics landscape, by offering sheer amounts of data about previously unaccessible domains in a cheap and scalable way. However, fast, reproducible, and cost-effective data analysis at such scale remains elusive. A key need for achieving it is being able to access and query the vast amount of publicly available data, specially so in the case of knowledge-intensive, semantically rich data: incredibly valuable information about proteins and their functions, genes, pathways, or all sort of biological knowledge encoded in ontologies remains scattered, semantically and physically fragmented.

**Methods and Results:** Guided by this, we have designed and developed Bio4j. It aims to offer a platform for the integration of semantically rich biological data using typed graph models. We have modeled and integrated most publicly available data linked with proteins into a set of interdependent graphs. Data querying is possible through a data model aware Domain Specific Language implemented in Java, letting the user write typed graph traversals over the integrated data. A ready to use cloud-based data distribution, based on the Titan graph database engine is provided; generic data import code can also be used for in-house deployment.

**Conclusion:** Bio4j represents a unique resource for the current Bioinformatician, providing at once a solution for several key problems: data integration; expressive, high performance data access; and a cost-effective scalable cloud deployment model.

## Introduction

### Big data in biology

In the past decade, Next Generation Sequencing in particular, and high-throughput technologies in general, have Read Archive [1] is growing exponentially, with 3.55445926862412 × 10^15^ bases at the time of this writing. More brought to Biology an exponential increase of what we could call raw data. The data available at the Sequence the UniProt TrEMBL database contains 9.2×10^7^ proteins [2]. And, without detracting anything from the importance recently, this wave is starting to be felt at the level of more structured, semantically-rich data: the latest release of of raw data, what is more and more a key need is having ways of not only storing raw data, but of also linking it with more structured knowledge, to be able to reason about it and connect it with other domains. That is where the focus of our work lies.

In the rest of this introduction we are going to describe the current challenges posed by big data in Biology, from the following dimensions:

- data characteristics
- data access
- data integration

### Data characteristics

We will first look at how this divide (admittedly spectrum) between raw and structured data could be defined, the relationships between the two kinds of data, and implications thereof.

Biological big data (inasmuch as is knowledge-intensive) tends to be big not only horizontally but vertically, exhibiting what we could call type-level complexity.

A good example of what happens in other domains could be Twitter data; there we can easily model data with a few vertex and edge types, while the typing restrictions on source and target vertices imposed by edges are fairly simple. What makes this challenging from a data perspective is the sheer number of values of each type.

In contrast, in the biological big data landscape there is a clear dichotomy between so to say “raw” and “structured”, semantically rich data. Sequencing reads from massively parallel technologies, or unprocessed data from other high throughput technologies such as microarrays or proteomics experiments would be clear examples of the first category. For the second we are thinking about, for example, data organized in ontologies such as functional information associated to proteins, genes and genomes, coming from organisms classified themselves in complex taxonomic structures.

Both kinds have become big, but across different dimensions: in the case of raw data it is the number of values of a given type, what we could call the *horizontal* complexity of data, that presents a challenge. In the other case, not only the data size but the sheer number of types and the intricate relationship between them (vertical complexity) is what makes them big. It is this second kind of structured, semantically complex data what we want to focus on, and to what Bio4j aspires to be a solution for its management and access.

The challenges and opportunities offered by these two different kinds of big data are also different, as dictated by their nature. In the case of raw data we are in a scenario which is not so dissimilar to what happens in other domains, such as high energy physics; however, in the case of structured data, the situation is unlike that present elsewhere. For example, just understanding a particular subfield requires years of training, and being able to (even partially) condense and incorporate that knowledge into the data model and query languages offers the chance of a breakthrough in how we work with this kind of data.

But this would impact not only structured data; there is of course a close, though highly asymmetric relationship between the raw and structured worlds. The access to raw data is normally mediated by previous filtering steps, where subsets of the raw data are defined using highly structured information linked with it. Furthermore, it could be argued that this does not happen more often in practice because of the absence of this link, and the lack of integration between different data sources. To put an example, it is not common to have the need of taking the sequence of all proteins ever known to men. What one normally wants is to get a particular subset determined by more high-level data such as protein function, taxonomy, interactions between them or their presence in certain processes, just to name a few possible conditions.

### Data access needs

Here we want to look at the requirements and peculiarities of accessing highly structured big data in the context of high throughput data analysis.

#### Accessing and querying highly structured data

Most data analysis processes need to deal with embarrassingly parallel computations requiring complex access patterns over highly structured (frequently immutable) big data. Obvious examples can be found in metagenomics studies, where in one way or another one needs to access taxonomies and data linked with them; or in comparative genomics, where we derive from the presence of genes and their encoded proteins in networks, the difference in biological function (encoded through different ontologies) between different genomes. Cancer genomics is another area where massive data analysis becomes challenging mostly because of the need to combine, in a complex way, many different elements and relationships. One needs to work with elements such as genomic variants, variant type, tumor type, tissue type, cell type, cellular compartment, gene type, protein motifs, protein interactions, enzymatic activities, complex functional aspects, and, most critically, relations between all of them. The importance of taking all these types into account is highlighted by, for example, the discovery of new therapeutic targets for personalized treatments based on monoclonal antibodies, or new biomarkers useful in diagnostics for some types of cancer, as exemplified by [3]. In these achievements being able to query data about genes and proteins and the intricate network of relationships between them has been crucial.

#### What, when, how

The temporal aspect is also key. Sometimes analysis can be scheduled (annotating a genome for a research project), while in other cases it is inherently impossible to even predict when you will need to run them (think of analyzing data coming from an outbreak). The resources needed or their characteristics vary wildly depending on the specific analysis. We are thus in a context where in general we do not know in advance what we need to access, how fast, when, and how.

It is in our opinion crucial guaranteeing that the big data explosion in biology in general, and genomics in particular does not bring as a consequence the confinement of data analysis basically to people or institutions with a lot of money and resources. The democratization of big data access (together with analysis tools) is definitely needed if we, as a community, want to be able to cope with the data deluge brought by NGS and related high-throughput technologies [as already identified in 4].

### Data Integration

As high throughput technologies extend their impact to more and more areas, the amount of domain-specific resources and data is inevitably going to grow. Of course linking them is (and will be) needed, but not **just** that; linking does not somehow magically integrates disparate resources and data models, it just puts the burden of integration on the user. As mentioned before, we need expressive ways of accessing structured data; but in most cases data is bound to come from different sources, with disparate data models.

It is important to stress that we are talking about the integration of two different data models through creating a new one which can act as a bridge between them; making the integration a first-class citizen makes possible the independent evolution of data models as well as the development of different integrations.

Integration is sometimes confused with data aggregation at the physical level; nothing could be further from the truth, as having a common bridge data model is a must for being able to query and access related but physically distributed resources.

We can speak of data integration at different levels; in decreasing order of abstraction

1. model – *what*
2. access and store – *how*
3. deployment and distribution – *where and when*

1 refers to what could be called the level of semantics: how we can reason about our data. Types, relationships, the dependencies between them, schemas; they all live here. In one word, this could be summarized as *what* is our data. In 2 we put the different ways of accessing our data, such as query languages, and related mechanisms and technologies for storing it; *how* we access our data. Finally 3 is concerned with the spatio-temporal aspects: *where* is the data (endpoints, deployments of infrastructure giving access to it) and *when* (availability, data as a service).

Changes in one level will obviously propagate to the next. Of course this layered classification is sketchy and imprecise, but we think that it can serve as a guide for discussing, categorizing and evaluating data integration efforts.

We want to stress that for integration to yield something useful, it should work at **all** 3 levels just mentioned: data integration efforts which only aggregate together different data models are imposing the burden of data model integration on the user (users actually, which invariably leads to a plethora of such integrations), with the added disadvantage of not being able to make any use of that integration at the other two levels.

## Materials & Methods

### Graph technologies

At the data storage and access levels, Bio4j is fundamentally based on a generic Java library for working with typed graphs, *Angulillos*, and Titan a scalable native graph database.

### Angulillos

*Angulillos* manuscript in preparation is a Java 8 library for defining and interacting with strongly typed graph data. Developed as part of Bio4j by the last author, motivated by the need of working with different graph data stores generically while taking advantage of all the information encoded in the Bio4j data model.

In Bio4j, *Angulillos* is used for both declaring the Bio4j data model at compile-time and as an embedded type-safe query language. This allows us to write data import code which is generic on the graph data store, and the user to write compile-time checked queries that can be executed transparently across different Bio4j distributions.

As of this writing, Bio4j uses *Angulillos* 

~~~
0.4.1
~~~

.

### Titan

Titan is an open source graph database developed by Aurelius with support for distributed operation, several storage backends, schema definitions, and local indices. In Bio4j, Titan is used for providing a Titan-optimized Bio4j distribution. As of this writing, Bio4j uses Titan 

~~~
0.5.2
~~~

, through an *Angulillos* API avaiable at bio4j/angulillostitan.

## Cloud Computing

### Amazon Web Services

Amazon Web Services (AWS) is the de-facto standard cloud computing platform, focusing on what is called IaaS (Infrastructure as a Service). It is composed of several services, some of them structured across different regions (US, Europe, etc) corresponding to the physical location of the resources provided. The two services on which Bio4j is based are EC2 and S3.

#### EC2

EC2 (Elastic Compute Cloud Service), part of the Amazon Web Services offering, provides the user with virtually unlimited horizontally scalable compute capacity, in the form of instances (virtual servers) of different types (hardware configurations and operating systems). The cost model is pay-as-you-use: AWS bills you per instance hour usage. You can provision an unlimited number of instances, while creation/destruction times are approximately 2 minutes.

We use EC2 as our basic compute infrastructure for all tasks, taking advantage of the service model, and the possibility of choosing hardware configurations on demand. Downloading the raw data from the different datasets is done using cheap standard instances, while the CPU-bound import process is done using instance types optimized for the task. Lastly, the ready-made configurations for deploying a Bio4j instance can use instance types with a considerable amount of RAM and fast SSD hard drives.

We interact with the EC2 service using the AWS SDK for Java, version (as of this writing) 

~~~
1.6.8
~~~

. We use the latest API version of the EC2 service itself, 

~~~
2013-10-15
~~~

#### S3

S3 (Simple Storage Service) is a horizontally scalable object storage system; you can store an unlimited number of objects (files), ranging from 

~~~
1B
~~~

 to 

~~~
1TB
~~~

 in size, inside a flat hierarchy of namespaces called “buckets” (folders). Objects are stored replicated across several zones, then across several data centers, then across several servers; the provided estimate for durability is 

~~~
99.999999999%
~~~

 per object per year.

In Bio4j, S3 is used for storing and distributing globally the data of each Bio4j release: objects correspond to the binaries of a Bio4j release already loaded into a particular graph database technology. We also cache the raw data coming from the different integrated datasets, so as to provide a globally available high-performance endpoint for their retrieval; transfer speed between S3 and EC2 is on the network attached storage level, thus reducing considerably import and release times.

We interact with the S3 service using the AWS SDK for Java, version (as of this writing) 

~~~
1.6.8
~~~

. We use the latest API version of the S3 service itself, 

~~~
2006-03-01
~~~

## Datasets

Bio4j integrates open data coming from different data sources; we include here a short description of each.

### UniProtKB

UniProtKB is the largest database of protein sequences and protein functional annotations freely available. It is composed by UniProt-SwissProt, which contains manually annotated and reviewed proteins, and UniProt-TREMBL, which contains automatically annotated and not reviewed entries. [5]

### Gene Ontology

GeneOntology (GO) is the main freely available database for gene ontologies. It is the major resource of controlled vocabulary for genes and proteins annotations. [6]

### UniRef

UniRef contains clusters of UniProtKB sequences. There are three kinds of clusters depending on the sequence similarity required to build the cluster: UniRef50, UniRef90 and UniRef100 where the members of the cluster share at least 50, 90 or 100 percentage of similarity respectively. [7]

### NCBI taxonomy

The NCBI taxonomy database is a publicly available curated collection of names and classifications for the organisms present at the GenBank database. [8]

### Expasy EnzymeDB

The Expasy ENZYME database is a collection of information and recommended nomenclature for enzymes. Enzymes are the kind of proteins responsible for catalyzing chemical reactions. It is the main publicly available resource for information on enzymes. [9]

### Bio4j codebase

The Bio4j codebase is open source, licensed under the open source approved AGPLv3 license. All development happens on GitHub, under the Bio4j organization.

## Results and Discussion

### Biological data, graph data

That most of the (particularly knowledge-based) data in biology comes in a graph-like form appears as something glaringly obvious. Just think of examples such as taxonomic or phylogenetic trees, signaling, metabolic or interaction networks, or the Gene Ontology. In most cases, data which we reason about using a graph model is forced to fit into a relational setting, creating, among other difficulties, a considerable impedance mismatch and far from optimal performance. But it is not only a question of effectiveness or complexity but of expressiveness: a semantically-guided analysis of the topological properties of such data has proven to be a source of important insights at the biological level. Some examples are

1. *Horizontal gene transfer detection* Searching for proteins in the same UniRef100 cluster but assigned to a different vertex of the taxonomy tree
2. *Protein motifs related with cancer and apoptosis* Searching for all the protein motifs (and their frequencies) in all genes annotated with some term related with apoptosis, requiring them to be members of a UniRef90 cluster in which there are proteins from different taxonomic vertices annotated with apoptosis terms.
3. *Proteins related with aging* First we select the species with the highest and the lowest life expectancy values. Then, we analyze the enzymatic activities that are exclusive of these two sets of species.
4. *Analysis of enzymatic activities and intelligence* We select a set with the most intelligent species and a control set of species. Then we analyze the enzymatic activities that are more abundant in the proteomes of the selected set with respect to the control.
5. *Analysis of GeneOntology terms in a set of differentially expressed genes* The GeneOntology terms from a set of differentially expressed genes can be easily retrieved from Bio4j; for a given set of GeneOntology terms a set of more meaningful set of GeneOntology terms related to the initial ones could be easily obtained traversing the Bio4j GO module. This kind of analysis (also known as GoSlim analysis) is of great help in, for example, the interpretation of RNAseq experiments, as they are used to infer the most representative functions in a set of genes.
6. *Analysis of the conservation of enzymatic activities* The conservation of enzymatic activities in a particular taxa could be measured using the size of the UniRef clusters for such activities (represented as EC numbers): bigger clusters imply higher conservation levels. Comparisons of the conservation degree among enzymatic activities or among taxa could be easily done.
7. *Characterization of functions encoded in plasmids* Plasmids are mobile genome elements found in Bacteria. They play a key role in important processes such as antibiotic resistance. The functions of the genes encoded in such plasmids could be easily obtained in Bio4j. Comparisons of these functions between pathogenic and non-pathogenic Bacteria could also be of interest, as a source of new insights into the functions responsible for their pathogenicity.

### The property graph data model

Note that in the examples above, we require a way of expressing properties about the graph structure itself together with typing restrictions, or predicates on vertices and edges; and this is exactly what the property graph data model offers. Property graphs have vertices and edges where both can have a set of properties; a property graph can be queried using so-called graph **traversals**, which return a subgraph based on the result of traversing the graph while matching edges and vertices with predicates over their types and properties. For the examples just mentioned, they could be translated to simple traversals over a convenient graph data model.

Due to the specific needs of Bio4j, we developed a strongly-typed generic version of it, called Angulillos. Thanks to it we can express the Bio4j graph model statically and write generic traversals that can then be run on any implementation of the Bio4j model.

### Bio4j dataset integration and graph data model

One of the motivations for the creation of Bio4j was the integration of the most important databases needed for bioinformatics analysis and especially for biological data interpretation. In the selection of data sources and the graph model design, a special interest was put in both the selection of data sources and the graph model design for Bio4j, keeping in mind how this could help for tasks that are cumbersome, difficult or almost impossible given the current state of APIs, integrations and access for the aforementioned resources.

Thus Bio4j includes protein data from UniProt, taxonomy data from NCBI and systematic annotation data from Gene Ontology, all of them composing a modular, truly integrated graph structure. By including all these databases we practically cover all the structured data needed for most of genomics, metagenomics, and transcriptomics analysis, for all species: from viruses to human.

The Bio4j data model was designed considering the intrinsic and extrinsic semantic features of its elements, the nature of their relationships and their importance from the point of view of biological knowledge extraction. Particular importance has been given to modeling connections between elements coming from different data sources, generally as edges going from one element type to another; this makes possible the extraction of conclusions involving different types of elements.

#### Integrated datasets

Here is the list of datasets integrated by the current Bio4j release, together with their relevance from a biological point of view.

#### UniProt

UniProt proteins are the crucial element in Bio4j; the data model puts proteins in the center as a hub of the network for the most important relationships. Another type of relationship between proteins included in Bio4j is the protein-protein interaction. The interactions between proteins annotated manually by UniProt curators were also modeled and incorporated to Bio4j. These connections make possible to draw networks of functionally related proteins. The complete UniProt dataset (SwissProt+TrEMBL) is included in Bio4j.

#### GeneOntology

Function prediction for sequences is the basic aim of bioinformatics, and to have standardized function annotations is crucial for obtaining biological meaning out of the sequence networks. GeneOntology (GO) is the major initiative to standardize the representation of gene and gene product attributes providing a controlled vocabulary of terms for their annotation. The modeling of the GO data was carried out maintaining the GO semantic and the directed acyclic graph (DAG) hierarchic structure of the GO data, but taking advantage of the possibilities of the graph paradigm since it is immensely easier to represent the relationships of the GO DAG in a a graph that in a table paradigm. The complex structure of tables that GO currently has, caused in a large degree by the relational database model in which it is implemented, precludes a flexible and efficient querying of the database. A graph though, is the natural structure to interact with GO elements and relationships. The integration of GO in Bio4j allows us to use a systematic annotation system including functional concepts that can be used to query the database for any of the elements and relationships included in Bio4j.

#### NCBI Taxonomy

Two of the main interests for researchers working with sequences are the organisms from which the sequences came from and the functions in which the sequences are involved. Bio4j includes the complete taxonomy tree from NCBI. UniProt taxonomy is also mapped to NCBI taxonomy, facilitating the access to functional data associated to proteins; such connections are incredibly useful in contexts such as the functional profile characterization of metagenomics samples. Indexes give you efficient access to the NCBI taxonomic vertices through GenBank Identifiers (GI), as is needed for example in the diversity analysis of genomic sequences.

#### UniRef

Considering that function inference for sequences is basically based on similarity we decided to include UniRef database in Bio4j. The UniRef database provides precomputed similarity analysis for the UniProt protein space; the availability of this information in the graph allows us to obtain functional data from the database with different levels of specificity.

#### EnzymeDB

As a prerequisite for the integration of metabolic networks data in Bio4j, data from EnzymeDB was modeled and included in Bio4j. The enzymes are the building blocks of the metabolic networks as they are the proteins responsible for the metabolic reactions. EnzymeDB database contains information on those enzymes characterized by the international EC number. For each enzyme in the database the following information is available: EC number, recommended and alternative name, catalytic activity and cofactors (if any)

Getting EnzymeDB data available and integrated in Bio4j as a graph facilitates the reconstruction of metabolic networks as the information is modeled in a more similar way as it really is. It also allows developing software for metabolic pathways analysis relying on complex queries and graph traversals.

#### Bio4j graphs

Following a modular approach, Bio4j is organized into different graphs. These graphs, as seen from the data sources perspective, can play two different roles: In one of them they correspond to a data source graph model (like 

~~~
UniProtGraph
~~~

, 

~~~
GoGraph
~~~

), including vertices and edges like 

~~~
protein
~~~

, 

~~~
proteinProteinIteraction
~~~

 or 

~~~
goTerm
~~~

, 

~~~
isPartOf
~~~

. But they also serve as graphs which add edges connecting data coming from different data sources; for example, there is a 

~~~
UniProt_go
~~~

 graph which adds and edge 

~~~
goAnnotation
~~~

 going from proteins to GO terms, representing all the GO terms with which a protein is annotated. Here we can see at play one of the advantages of graph data models: so-called cross-references can be treated as just another type of data, modeled as edges connecting existing vertices.

Of course, it would not make sense to have 

~~~
UniProt_go
~~~

 if you do not already have both the 

~~~
UniProt
~~~

 and the 

~~~
Go
~~~

 graphs; graphs have *dependencies*. This is taken into account in our model, making it possible to simultaneously achieve modularity and correctness: the user is free to use the graphs that he may needed as long as he includes the corresponding dependencies.

#### The data model:highlights

The Bio4j data model is too big be thoroughly described here; instead we offer a set of samples from different data sources, highlighting how the typed graphs approach can help.

#### GeneOntology

In this case the native data model is really close: the GeneOntology (GO) is a directed acyclic graph, with typed vertices and edges; edges are the ontology relationships (*is part of*, *regulates*, etc) while there is essentially just one type of vertex: a *term*. The typed graph approach also shines here when modeling GO domains: we do not need to modify terms here, just add a *domain* vertex type and add “witness” vertices for each of the three domains (*cellular component*, *biological process*, and *molecular function*) together with edges going to each of the terms. These edges work as typing ascriptions, while making possible the independent evolution of the GO domains structure: for example, adding / deleting new domains or lifting the restriction of each term living in exactly one domain is straightforward, and most importantly does not involve modifying terms in any way.

#### NCBI taxonomy

The modeling of the NCBI taxonomy in Bio4j is pretty straightforward as well. The NCBI taxonomy is a tree where the vertices are *taxa* and the edges are the phylogenetic relations among taxa (mainly a child vertex *is a* parent vertex).

The protein taxonomy assignments were modeled as edges between the proteins vertices and their corresponding taxa vertices.

#### Protein relationships in UniProt

Relationships between different proteins such as protein-protein interactions are modeled as edges between them. This makes possible, for example, the analysis of the topology of the corresponding networks, such as shortest paths between proteins, connected components, or the determination of clusters according to complex criteria. The integration of other resources such as GO annotations adds another dimension to this, in that a topological analysis can be complemented with functional information.

## Bio4j deployment, distribution and cloud computing

As mentioned in the Introduction, the Bioinformatics high throughput data analysis space has a set of really specific characteristics and requirements; we strongly believe that they match perfectly with the cloud computing and data as a service paradigms. Bio4j was designed from the start with this in mind.

### Using Bio4j as a service

A full import of all the integrated data sources is stored in a public S3 bucket, which allows anyone to create an EC2 instance and use Bio4j in 2 minutes. Data as a service in this sense brings the democratization of big data studies; having all the data from critical resources such as UniProt, UniRef, or Gene Ontology readily integrated and wrapped in a highly tuned graph database, at the cost of grabbing a coffee from the coffee machine represents a dramatic paradigm shift.

There are also obvious benefits in terms of reproducibility and data exchange, not to mention performance: you can deploy hundreds of copies of Bio4j giving high-speed local access and workflow-specific parallelization at a reasonable cost.

### Bio4j on your own infrastructure

The import code is completely generic and will run on any environment where Java 8 is available. If the user does not want (or simply cannot) use Amazon Web Services, he can download the raw data and run the import code on his own infrastructure.

## Conclusions and future work

Bio4j allows extracting biologically sound conclusions difficult to obtain with the current state of biological databases. It demonstrates that a typed graph model is a perfect fit for storing, querying and reasoning about biological data.

### Future work

The integration of other publicly available data sources such as Reactome [10] is already underway; we plan to add network data from other freely available databases in future Bio4j releases. The other key data resources which we plan to integrate in the following months center around genomics data [11], where a graph model can help in (among other things) modeling and querying position-based data such as annotations.

On the API side, we are working on a new Scala version, together with improvements in the cloud Bio4j distribution.

## Acknowledgments

We wish to thank

- The **Google Summer of Code 2014 program**, where Bio4j was a participant as a mentoring organization.
- **Alexey Alekhin** and **Evdokim Kovach** for helpful discussions and their involvement in related developments around Bio4j.

Bio4j has been partly funded by the INNPACTO program grant IPT-2011-0735-010000.

